# Risks to pollinators from different land-use transitions: bee species’ responses to agricultural expansion show strong phylogenetic signal

**DOI:** 10.1101/524546

**Authors:** Adriana De Palma, Michael Kuhlmann, William D. Pearse, Emma Flynn, Stuart P.M. Roberts, Simon G. Potts, Andy Purvis

## Abstract

Bee species worldwide are facing a future of further land-use change and intensification. Populations of closely-related species with similar ecological characteristics are likely to respond similarly to such pressures. Such phylogenetic signal in species’ responses could undermine the stability of pollination services in agricultural and natural systems. We use abundance data from a global compilation of bee assemblages in different land uses to assess the sensitivity of 573 bee species to agricultural expansion, intensification and urbanization; and combine the results with the Bee Tree of Life to assess phylogenetic signal. In addition, we assess whether variation in species’ sensitivity to land-use change is better explained by phylogenetic or available functional trait differences. Bee species show strong phylogenetic signal in sensitivity to agricultural land expansion but only a weak signal in sensitivity to agricultural intensification and urbanisation. Sensitivities were usually best explained by a combination of functional and phylogenetic distances. This finding suggests that the commonly-recorded traits, despite being meaningful as functional response traits, do not capture all important determinants of bee species’ vulnerability or resistance. However, it also suggests that model-based predictions of the sensitivity of poorly known species may be sufficient to help guide conservation efforts.

## Introduction

Land-use change is the most important driver of present-day terrestrial biodiversity loss [1,2] and is predicted to cause continued damage in the future [3,4]. Most scenarios incorporate further loss of natural and semi-natural land, driven by agricultural expansion and urbanisation, as well as increased degradation due to agricultural intensification [5]. The effect such global change will have on bee populations could have serious consequences for crop pollination worldwide [6] and— as bee species are the most important pollinators of flowering plants globally [7]—for wild plant populations too [8].

Predicting likely effects of land-use change on ecosystem functions such as pollination requires an understanding of whether and how species’ sensitivities to land-use change are shaped by their functional traits and evolutionary history [9,10]. Traits are known to predict responses of European bee populations to land use and related pressures, but associations are inconsistent [11–13]. In bumblebees, long-term population declines in North America show phylogenetic signal (i.e., closer relatives tend to show more similar trends) [14], as does their global conservation status as assessed by the IUCN Red List [15], but information on other bees is limited. The relative usefulness of phylogeny and available functional trait data in predicting species’ responses is an open question. Although phylogenetic relatedness is in this context only a proxy for similarity of species’ functional responses [16], it may outperform functional trait data if responses are driven by a broader set of (phylogenetically patterned) ecological differences than are captured by available trait data.

A strong phylogenetic signal in species’ responses to a particular pressure increases the risk that the pressure could impact ecosystem functioning [16], as the order of species losses can strongly influence pollination networks [17]. Phylogenetic diversity and redundancy of bee communities may also decline; although debate is ongoing [18,19], phylogenetic diversity can be important for ecosystem functioning and stability [20]. The protection of phylogenetic diversity (by conserving those clades that are most vulnerable) may also be important for maintaining robust pollination networks: high phylogenetic diversity can correlate with interaction diversity [21], as closely-related species also tend to share resources [22,23]. The strength of phylogenetic signal can vary among pressures [e.g. 24], meaning that different land-use transitions could carry different risks of pollination impairment that might not be apparent from the effects on species diversity or overall abundance. Nonetheless, so far there has been no exploration of the phylogenetic pattern of bee sensitivities to particular threats such as agricultural expansion, agricultural intensification and urbanization.

Understanding which disturbances prompt the most phylogenetically patterned response may therefore indicate where ecosystem services might be at greatest risk and so where conservation action will be most important. However, it is unclear whether responses to different pressures will all show phylogenetic signal [16] as the links between ecological traits and the response to human impacts are not always straightforward [11,16]. We use data from 86 studies and 2,599 sites to assess the sensitivity of 573 bee species (from 96 genera) to human-dominated land uses, including agriculture and urban areas, and to increasing agricultural intensity. We quantify the strength of phylogenetic signal in these sensitivity estimates, accounting for uncertainty in sensitivities. Using trait data for a subset of 143 bee species, we assess whether phylogenetic differences are better able than functional traits to explain variation in species’ sensitivities.

## Materials and methods

Site-level data on bee abundance and occurrence were extracted from the PREDICTS database [25] and De Palma et al. [26]. We refer to each survey of multiple sites that used the same sampling method within the same season and the same country as a ‘study’. Differences in sampling effort among sites within a study were corrected for where necessary, assuming that recorded abundance increases linearly with sampling effort [26]. Within each study, we recorded any blocked or split-plot design. See Appendix Table S1.1 for a list of data sources (some containing multiple studies).

The major land use and land-use intensity at each site was recorded based on information in the associated paper, supplementary files or other information provided by the original authors, as described in [27]. Land use was classified as primary vegetation, secondary vegetation, cropland, plantation forest, pasture or urban. The use-intensity scale assesses human disturbance on a three-level qualitative scale within each land use (minimal, light and intense) [28]. For instance, intensively-used cropland includes monocultures with characteristic features of intensification (e.g., large fields, high levels of external inputs, irrigation and mechanization); lightly-used cropland would show some, but not all, or the same features; whereas minimally-used cropland would include small mixed-cropping fields with little or no external input, irrigation or mechanization.

### Species sensitivity

We focus on four land-use transitions (Table 1). The first, with the largest sample size, compares species abundances in natural/semi-natural land (i.e., primary or secondary vegetation) with all human-dominated land uses combined (i.e., all other land-use classes) as a recent synthesis showed that, in terms of species composition, assemblages in human-dominated land uses are more similar to each other than to those in natural or semi-natural land [3]. However, within this broad categorisation, particular transitions may influence species in different ways, so we also explore separately the impact of conversion to agricultural land, conversion to urban land, and increases in agricultural intensity. The land-use classes within the dataset were coarsened depending upon the land-use comparison of interest, allowing species abundances to be compared between intact (natural and semi-natural) land uses and converted (human-dominated, agricultural or urban, respectively) land uses, or with increasing agricultural intensity (Table 1).

**Table 1:** Land-use transitions of interest and the component land-use classes. Dataset sample sizes are also given: number of studies, species and species that match with the recently-published Bee Tree of Life [32].

| Land use comparison | Intact | Converted | Number of studies | Number of species | Number of species in the phylogeny |
| --- | --- | --- | --- | --- | --- |
| Loss of natural and semi-natural habitat | Primary & Secondary vegetation | Cropland, Pasture, Plantation forest & Urban | 86 | 488 | 141 |
| Agricultural expansion | Primary & Secondary vegetation | Cropland, Pasture & Plantation forest | 84 | 437 | 121 |
| Urbanisation | Primary & Secondary vegetation | Urban | 54 | 146 | 56 |
| Agricultural intensification | Agricultural sites (as defined above) with a continuous use-intensity scale from 1 (minimal use) to 3 (intense use) |  | 53 | 268 | 87 |

We define the Species Sensitivity Index (SSI) as the log-response ratio:

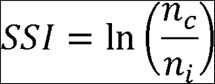

where *n*_*c*_ is the mean abundance of the species in converted land uses, and *n*_*i*_ is the mean abundance in intact land-uses. SSI was estimated as follows. For each species, abundance was modelled statistically as a function of the land-use category (intact vs converted) or agricultural intensity. Study identity was included as a factor to account for among-study differences in methodology, sampling effort and biogeography. Study identity would ideally be treated as a random variable, but many species were in too few studies to accurately estimate random-effect variances [29] (~80% of species in our dataset were represented in six or fewer studies). Bayesian generalized linear models were used [with weakly informative default priors, as described in 30, implemented in the arm package using the bayesglm function] with a Poisson error structure unless overdispersion required use of a quasi-Poisson structure [31]. This approach provides more realistic coefficient estimates and more conservative standard errors than frequentist generalized linear models given that there can be complete separation of the data (for example, when a species is always absent from converted sites but always present in intact sites). The land-use coefficient (the log response ratio) from the model was used as the estimate of SSI and the standard error as its associated uncertainty.

In all further analyses, we use data for species that have abundance values in at least six sites (three intact and three converted), aiming for a balance between maximizing numbers of species and having sufficient data for each one. Repeating the analyses with a more stringent threshold (at least 12 sites: six intact and six converted site) produced qualitatively similar results, so we show results with the more lenient threshold, which include more species.

### Phylogenetic tree

We used the Bee Tree of Life [32], a recent phylogeny of over 1,300 bee species from around the world, after rate-smoothing using PATHd8 [a computationally efficient method for large phylogenetic trees 33] with the root age constrained to one. Of the species present in our dataset, 141 were also in the phylogeny. Although there was no difference in mean SSI between these and the species absent from the phylogeny (two-tailed t-test: t_1,245.0_ = −0.33, n.s.), the latter tended to have higher uncertainty in SSI (two tailed t-test: t_1,230.1_ = −4.29, *p* < 0.001). Because of this non-randomness, we used the *pastis* package in R [a birth-death polytomy resolver, 34] to estimate—1000 times—possible placements for missing species, given their taxonomic affinities [35], to produce 1000 complete trees. See Appendix S2 for full details. Birth-death polytomy resolvers can bias trait-based analyses [36], so we perform all analyses on the rate-smoothed incomplete tree of 141 species as well as the 1000 complete trees.

### Phylogenetic analyses

Phylogenetic signal was quantified using Pagel’s [37] λ, which produces reliable estimates when sample sizes are large [38], as here. We used the implementation in the R package *phytools* [39,40], which accommodates uncertainty in the SSI estimates, though we also estimated λ without accounting for uncertainty in species sensitivity, for λ comparison. As bumblebees (*Bombus*) may respond differently from other species to human impacts [14,41,42], which could drive phylogenetic signal in the overall dataset, we also estimated λ for both bumblebees alone and all other species.

We used ‘traitgrams’ [43] to assess the relative power of phylogenetic and functional distances to explain variation in SSI. Phylogenetic and functional distances were combined into a single set of Euclidean distances [43] according to:

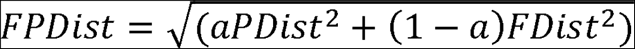

where *PDist* is the phylogenetic (cophenetic) species distance [*pez* package: 47], *FDist* is the trait pairwise distance [Gower’s dissimmilarity with transformation to provide Eucleidian properties: 48,49], and *a* governs the relative weighting of *FDist* and *PDist* in *FDist*. Setting *a* = 0 computes functional diversity and *a* = 1 computes phylogenetic diversity. We calculated 11 distance matrices using *a* values spaced evenly from 0 to 1, and compared their explanatory power to find which best explained our data in distance-based generalised linear models [dbstats package, 47,48] weighted by the inverse squared standard errors of the SSI. We also modelled SSI as a function of a randomly-generated distance matrix (with original values drawn from a normal distribution) as a null model. Models were compared using an adapted version of Akaike’s Information Criterion (AIC) [47,48]. We calculated models’ pseudo *R*^2^ as: 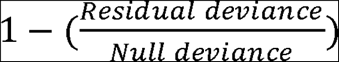

Ecological trait data were collated for 556 of the species in our dataset by SPMR, MK and EF by assessing peer-reviewed and grey literature and by measuring available museum specimens. Traits included inter-tegular distance (ITD) [a proxy for foraging range, 52], flight season duration, dietary breadth, nesting strategy, and reproductive phenology and strategy [11], and are generally phylogenetically conserved (see Appendix Table S3.2). However, because data for the complete set of seven traits were available for only 75 species in both our data set and the original phylogeny, we re-ran this analysis using a reduced set of traits (ITD, nesting strategy and reproductive strategy), enabling inclusion of 143 species. We also repeated the analysis using the completed trees (259 species with data for all traits; 537 species with the minimal set).

## Results

### SSI values

Species’ sensitivities to different land-use transitions were relatively normally distributed (see Fig 1), with wide variation in species responses. For the comparison between natural and human-dominated land-uses, for instance, 40 species had SSI < − 2 and 28 species had SSI > 2; 82 species had SSI estimates that were significantly non-zero (44 negative, 38 positive; coloured lines on Fig 1).

**Figure 1:**
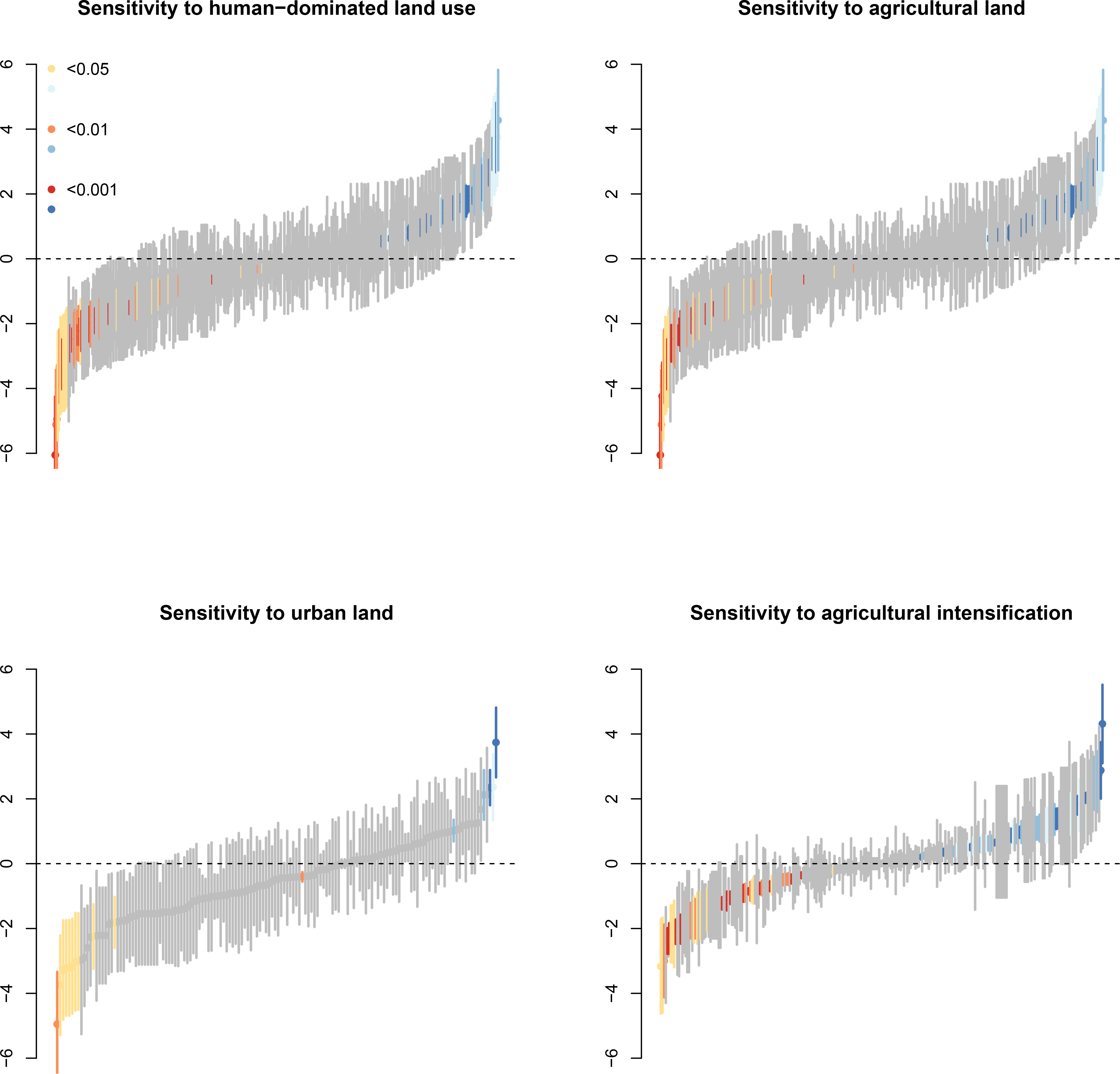
Spread of SSI values (points) and their standard error (lines) for each land-use transition, for all species in the dataset. Coloured points indicate that SSI values were significantly different from 0. Particularly vulnerable species (i.e., those with an SSI significantly below 0) are coloured in yellow to red, while species that strongly benefit from land-use change (i.e., those with SSI values significantly above 0) are coloured light to dark blue.

### Phylogenetic signal

The strength of phylogenetic signal in SSI differed among land-use comparisons (Fig. 2a). The strongest signal was in comparisons between (semi-)natural and human-dominated land uses, with high and significantly non-zero λ values for all but one of the completed phylogenetic trees (*p* < 0.05; Fig. 2a and Fig. 3). SSIs comparing semi-natural and natural land to agriculture showed λ values nearly as high: λ was significantly non-zero for the rate-smoothed tree (λ = 0.59, *p* < 0.05), and for >99% of the completed trees. The signal in sensitivity to urbanization and to increasing agricultural intensity depended on which phylogeny was used, being very low except with the completed trees (Fig. 2a). Weighting the analyses by the standard error of species sensitivity was extremely important: λ values were always low when standard errors were not accounted for (Fig. 2b).

**Figure 2:**
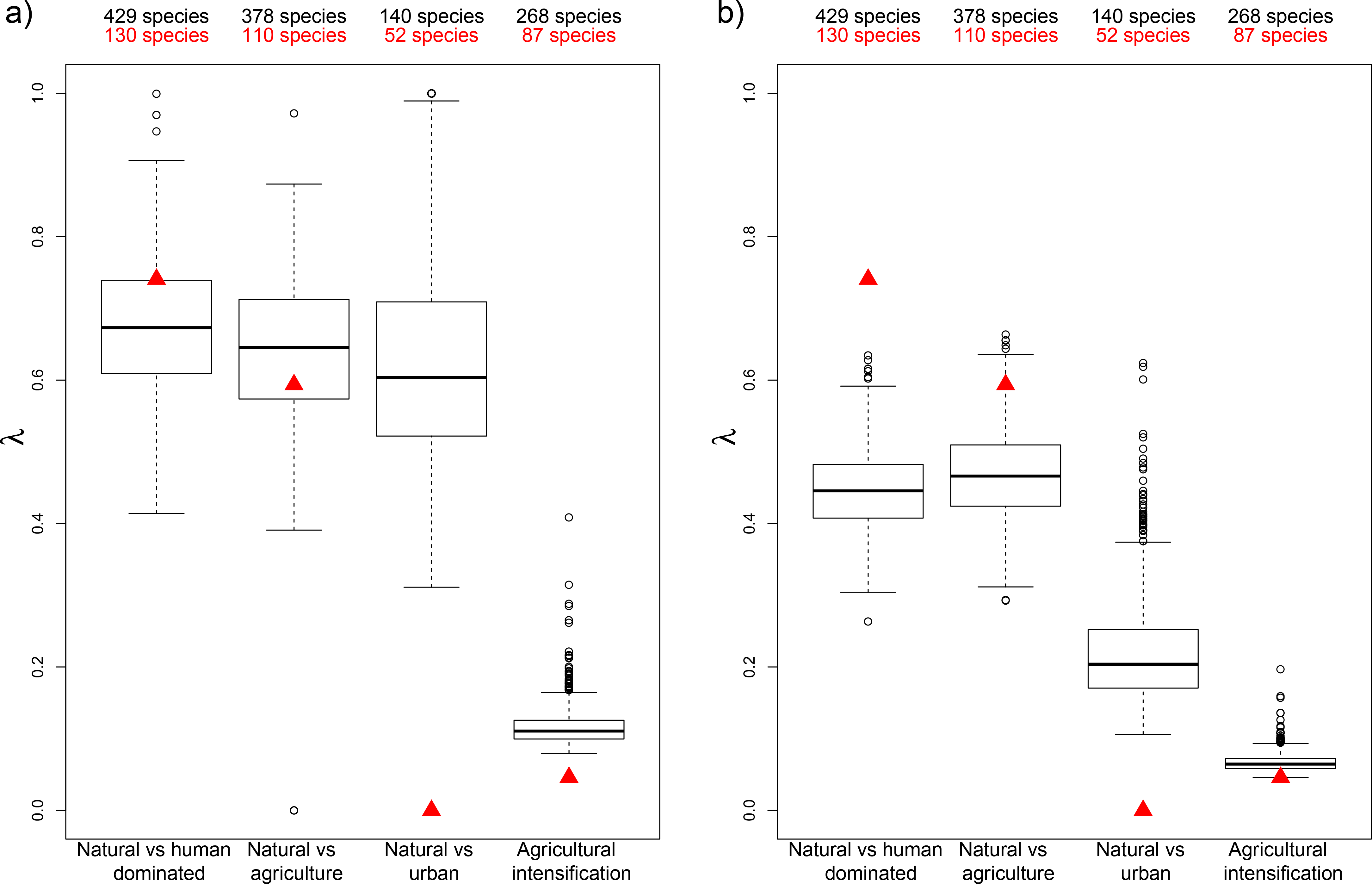
Phylogenetic signal in species’ sensitivity to various land use pressures, when (a) standard errors are accounted for and (b) when they are ignored. Red triangles indicate the values for the rate-smoothed tree, where some species are missing from the tree. Numbers in red show the number of species included in these tests. Boxplots show the distribution of values across the 1,000 completed trees, with numbers in black showing the number of species included in these tests.

**Figure 3:**
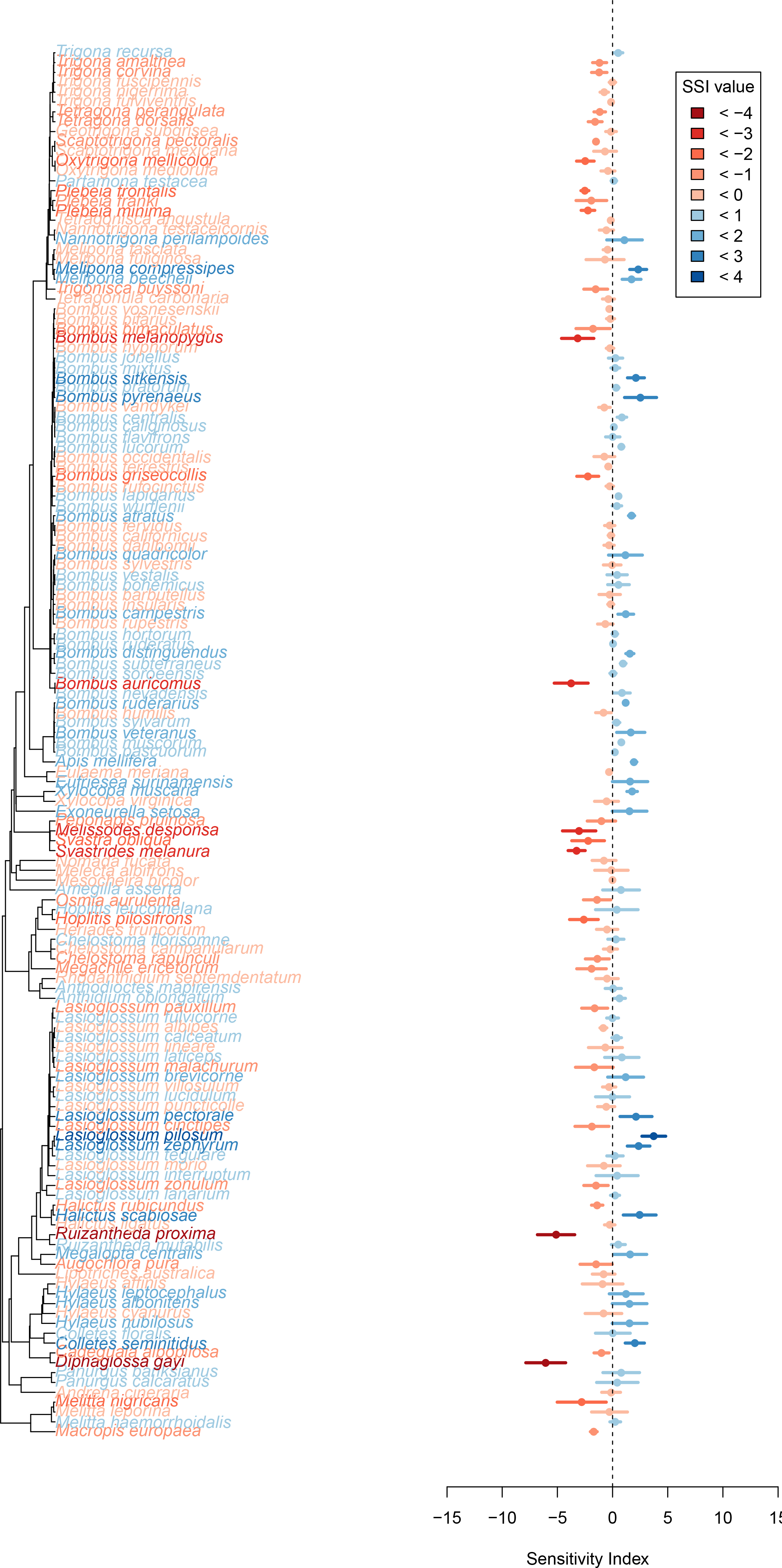
Phylogenetic signal in species’ sensitivity to all human-dominated land uses using the rate-smoothed (incomplete) tree. Tips of the phylogeny are coloured according to the species’ sensitivity: from blue to red indicate less to more sensitive. The right panel shows species sensitivity ± the standard error. See Appendix Figures S4.2 to S4.4 for similar figures for other land-use transitions.

There was no significant phylogenetic signal within the bumblebees. Phylogenetic signal was also lower among non-bumblebees, particularly when assessing species’ sensitivity to all human-dominated land-uses, and particularly when analyzing only the species in the rate-smoothed tree (Table 2); analyses using the complete phylogenies often found significant, albeit reduced, phylogenetic signal in non-bumblebees for both agricultural expansion and loss of natural vegetation (Table 2).

**Table 2:** λ values of the sensitivity of all bee species, bumblebees only and other species (bumblebees excluded from the phylogenetic tree) to land-use change. For the completed trees, this is the mean λ ± standard deviation across the 1000 phylogenies.

| Sensitivity to: | Tree | $\lambda$ | | |
| --- | --- | --- | --- | --- |
|  |  | All species | Bumblebees | Other species |
| Human-dominated land | Rate-smoothed | 0.741 | 0 | 0 |
| | Completed | 0.672 ± 0.09 | 0 ± 0 | 0.626 ± 0.14 |
| Agricultural land | Rate-smoothed | 0.594 | 0 | 0 |
| | Completed | 0.64 ± 0.1 | 0 ± 0 | 0.586 ± 0.16 |
| Urban land | Rate-smoothed | 0 | 0 | 0.143 |
| | Completed | 0.633 ± 0.15 | 0.256 ± 0.44 | 0.2 ± 0.12 |
| Higher-intensity agriculture | Rate-smoothed | 0.046 | 0 | 0.041 |
| | Completed | 0.116 ± 0.03 | 0 ± 0 | 0.121 ± 0.04 |

### Relative importance of traits and phylogeny

Species’ sensitivities to human-dominated land uses were best explained by a combination of both phylogeny and functional distances: this was true when using the rate-smoothed tree and across the completed trees (with α ≥ 0.7 producing the highest AIC weights and high R^2^ values; Figure 4 and Appendix Figure S5.1). However, phylogenetic distance was more important than functional distances for transitions whose SSIs showed higher phylogenetic signal (i.e., sensitivity to human dominated land and agricultural land), whereas sensitivity to agricultural intensification seemed to be more strongly influenced by functional distances (α < 0.4 had the highest AIC weights and R^2^ values; Figure 4 and Appendix Figure S5.1).

**Figure 4:**
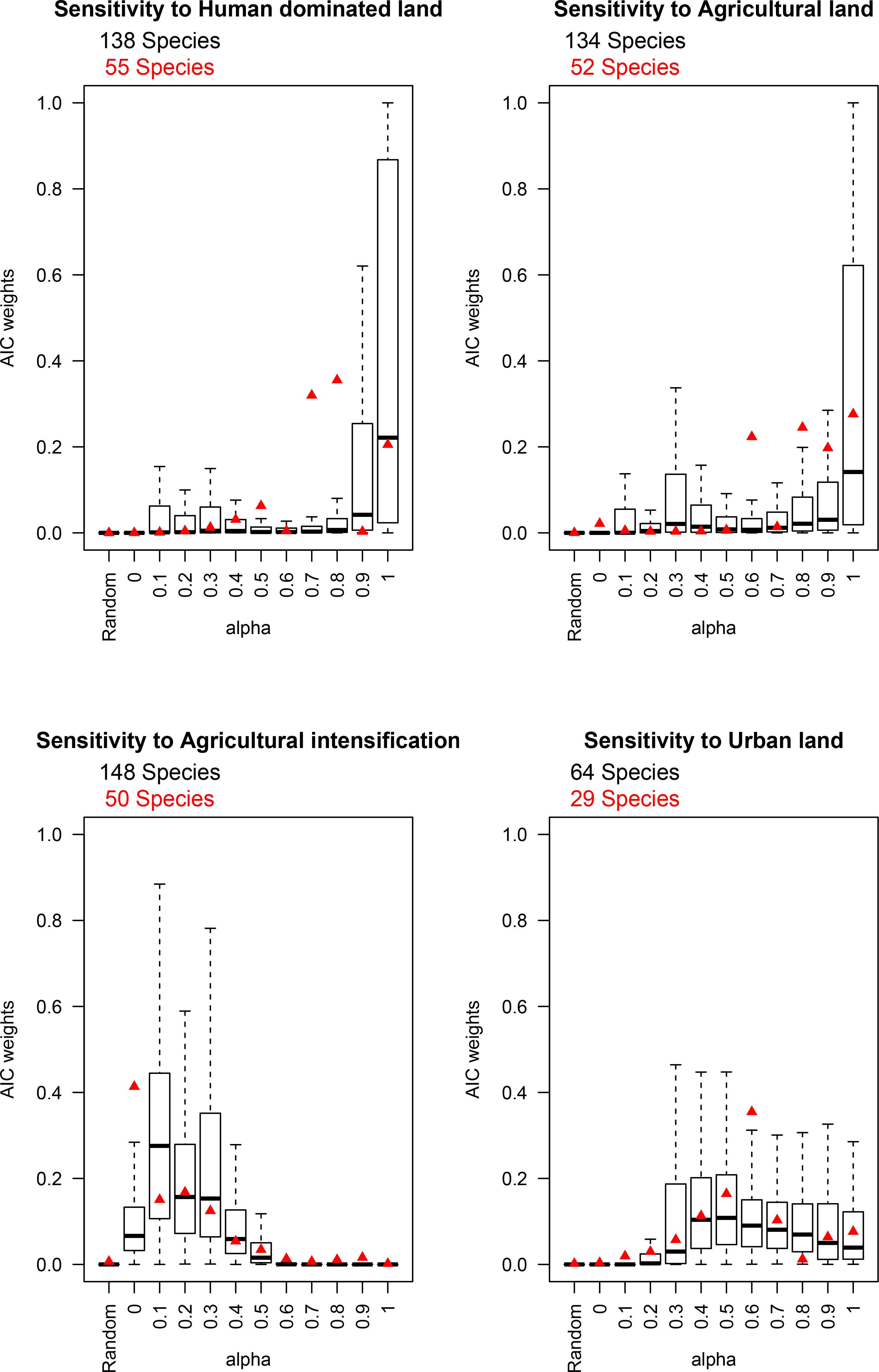
Akaike’s Weights of models assessing species sensitivities as a function of *alpha* value, where *alpha* = 0 uses only functional traits to calculate the distance, *alpha* = 1 uses only phylogeny to calculate the distance, and *alpha* = 0.5 considers both traits and phylogeny equally. Model performance was also calculated where distances were randomly computed. Red triangles indicate the values for the rate-smoothed tree, where some species are missing from the tree. Numbers in red show the number of species included in these tests. Boxplots show the distribution of values across the 1,000 completed trees, with numbers in black showing the number of species included in these tests. For a similar figure assessing explanatory power of models, see Appendix Figure S5.5.

## Discussion

Bee species show strong heterogeneity in their responses to land-use practices with, in our data set, roughly as many ‘winners’ (positive SSI) as ‘losers’ (negative SSI) (Fig 1). This is in line with the wide range of responses reported in the literature (e.g., urbanization has been reported as having positive [50,51], neutral [52] and negative [50,51,53] effects on bee populations). Clearly many bee species are able to benefit from the environmental changes (potentially including removal of competitors: [54]), while others struggle to persist.

The strong phylogenetic signal (λ ~ 0.7) seen in species’ responses to human-dominated land uses is not surprising: SSI estimates a species’ susceptibility to one particular driver (land-use change), which will be mediated by functional response traits [11,12,55,56] that are often phylogenetically conserved [57] (though estimated responses will also be influenced by the contexts of individual studies, probably reducing the phylogenetic signal).

The different focused land-use comparisons find different degrees of phylogenetic signal in species’ responses (Fig 2), with conversion to agriculture and urbanization producing moderately strong signal (λ ~ 0.6, except for the analysis of urbanization using the rate-smoothed tree, where sample size is small) while agricultural intensification elicits negligible signal (λ ~ 0). Accounting for measurement error was crucial for estimating phylogenetic signal strength; omitting it led to much lower estimates of λ (Fig 2), as expected [39], though results were consistent. Results were also not strongly driven by bumblebees; clustering of sensitivity was seen throughout the tree.

Why does agricultural intensification not produce a stronger phylogenetic signal? We consider three possible biological explanations. First, species’ responses to agricultural intensity may be mediated by features of landscapes—not considered in our models—more than by features of sites. Field diversity and landscape composition, for example, have been shown to shape the biotic effects of agricultural intensity [58]; because such attributes are not recorded in the PREDICTS database, we were not able to consider them, perhaps causing responses to appear idiosyncratic. Second, the low-intensity agricultural habitats to which higher-intensity agriculture is compared may have already filtered out the most vulnerable native bee species (i.e., those with the most negative SSI values), and perhaps had synanthropic (i.e., SSI > 0) non-natives added, by the original conversion from natural to agricultural land. Such SSI-biased turnover of species would tend to erode SSI’s phylogenetic signal. This explanation—that the initial conversion of a landscape to an ‘anthrome’ [59] leaves a strong phylogenetic signal whereas subsequent intensification of use does not—requires that much the same traits mediate responses to agricultural expansion and agricultural intensification. Such trait-based extinction filters are likely why mammalian extinction risk shows no phylogenetic signal in regions with a long history of intense human activity [60]. If this explanation is correct, then even though low-intensity agricultural practices score better than more intensive systems for community-level measures of bee biodiversity [35,58], they may have already altered community composition irreversibly [42]. The third possibility is that the ability of species to persist in the face of agricultural intensification depends most strongly on traits that are not phylogenetically conserved. For instance, we found no strong phylogenetic signal in diet breadth (Appendix Table 3.2). Diet breadth can be flexible for some species (e.g. *Bombus terrestris* can increase diet breadth when faced with increased resource competition [61]), but on the other hand, even generalist species can have rigid host-plant preferences [62]. A fourth, non-biological, explanation for the lack of phylogenetic signal in responses to agricultural intensification is that our use-intensity criteria mix multiple pressures (e.g. pesticide and fertilizer use); each component might on its own elicit phylogenetically patterned responses, but the signal may be lost by mixing them.

Although SSI values usually showed phylogenetic signal, in the subset of bee species with available trait information, SSIs were generally better explained by a combination of both phylogenetic and trait distances [43], highlighting the added value of ecological trait data for species. These results are in line with previous analyses revealing that functional traits can significantly influence species sensitivities to a number of land uses and land-use practices in a variety of systems [11,12,55]. However, such trait-environment relationships can have low explanatory power and results across studies can be contradictory [13]. Taken with our results, this suggests that species’ sensitivities may be influenced by ecological differences that are not fully captured by the commonly measured traits, while phylogeny may provide a closer approximation to these unmeasured characteristics [63]. For instance, the phylogenetic relatedness of bee species can inform the structure of plant-pollinator networks [23,64]; as ecological interactions are lost more quickly than species from a system [65], a given species’ network could have a strong influence on its resilience to disturbances. Responses to agricultural intensification do not fit this model: trait differences explained some variation in SSI but phylogenetic distance did not. This combination of results is consistent with the suggestion that trait-mediated competition may underlie the responses [66].

Although the importance of traits in mediating species’ sensitivities to land-use changes is congruent with previous work [12], our dataset may not be representative; it is therefore possible that a more complete set of species with trait data would change the relative importance of traits and phylogeny. The species in our dataset capture significantly less phylogenetic diversity than expected from a random selection of species from the Bee Tree of Life (see Appendix 6). Local assemblages are often a non-random subset of the global phylogeny [67,68]; our analyses focus on such assemblages and so this is, in part, an inescapable consequence of our study’s objectives.

The strong phylogenetic signal in bee species’ sensitivities to human-dominated land uses like agriculture means that losses of diversity are likely to be concentrated within a subset of clades, where they will be correspondingly more severe; likewise, any gains in diversity will be restricted to groups of related species. Clustering of losses can greatly reduce the phylogenetic and functional diversity of a system [16], potentially jeopardizing its ability to function under environmental change [20]. Crop pollination may be more robust than other ecosystem functions to species losses as a few dominant species are the main contributors at local scales [69]. However, higher species diversity may be necessary at larger spatial scales [70]. Furthermore, some wild plant species require specialist pollinators, potentially meaning that they face a double threat from conversion of land to agriculture: directly, though loss of habitat, and indirectly, through the decline in their pollinators.

Our results suggest that the phylogenetic risk assessment framework set out by Díaz et al. [16] could inform management practices and highlight gaps in knowledge. Even though bees are well studied relative to many other invertebrate groups, there are still uncertainties about their current status and vulnerability to human impacts [71] as well as gaps in trait data. The strong phylogenetic patterning in species’ sensitivities to agricultural expansion could help to predict the sensitivity of understudied species, identifying those that are most vulnerable or resistant to guide conservation planning [72,73]. For instance, many species of bumblebees showed positive or neutral responses to agricultural land, but negative responses to increased agricultural intensity (Appendix Figure S4.2 and S4.4); this combination of results suggests that agricultural production can support many species of bumblebees, but only if intensity is low. However, our estimate of sensitivity to particular pressures—SSI—does not necessarily indicate the conservation status of a species, which is a product of both sensitivity to combined pressures and exposure to those pressures. This may explain why we found no phylogenetic signal in bumblebee SSIs, even though extinction risk in these species is significantly non-random [15]. Phylogenetically patterned responses may also open opportunities to monitor bee communities at higher taxonomic levels [74–76]: this would reduce the need for species identification by expert taxonomists and the need for destructive sampling of bees [77,78], save time and money, and facilitate citizen science.

Our results provide a basis from which to make, test, and inform predictions about bee species’ sensitivities to land-use change, with potentially important benefits for monitoring and conservation prioritization, as well as identifying land-use pressures that may most affect pollination services to crops and wild species. Analyses of how these results scale up to changes in abundance-weighted phylogenetic diversity of communities are necessary to identify spatial patterns in diversity and potential areas of pollination deficit [35,79].

## Acknowledgements

We are grateful to all those who contributed data to this project, particularly members of the Greenveins project, and the PREDICTS team for assistance with data collation and curation, including Sara Contu, Sam Hill, Helen Phillips, Tim Newbold and Lawrence Hudson. We thank Gavin Thomas and Susy Echeverria-Londoño for assistance with phylogeny estimation. Much of the trait data used in this work was compiled by the EU FP7 project “Status and Trends of European Pollinators” (244 090, www.STEP-project.net). This work was supported by the BBSRC (grant BB/F017324/1 to ADP) and NERC (grant NE/M014533/1 to AP) and is a contribution from the Imperial College Grand Challenges in Ecosystems and the Environment Initiative. PREDICTS is endorsed by the GEO-BON.

## Author Contributions

ADP and AP conceived the study. ADP carried out statistical analyses and wrote the first draft of the manuscript. MK assessed species taxonomy. EF, MK, SPMR and SP collated trait data. WP provided guidance on phylogenetic analyses. All authors contributed significantly to revisions of the manuscript. All authors gave final approval for publication.

