## Appendix for "Risks to pollinators from different land-use transitions: bee species’ responses to agricultural expansion show strong phylogenetic signal"

**Appendix: Varying phylogenetic signal in bee species’ sensitivity to different land uses**

**Appendix S1: Diversity data**

**Table S1.1: Sources used to calculate species sensitivity indices**.  †Data are accessible via the PREDICTS database (which will be made openly available). ‡Data are available from the referenced paper. For all other datasets, please contact the corresponding author of that paper directly. See De Palma et al. (De Palma et al. 2016) for further summary data regarding these sources.

| **Source** | **Reference** |
| --- | --- |
| Kruess & Tscharntke | Kruess, A. & Tscharntke, T. Grazing intensity and the diversity of grasshoppers, butterflies, and trap-nesting bees and wasps. *Conservation Biology* **16**, 1570–1580 (2002). |
| Vázquez & Simberloff | Vázquez, D. P. & Simberloff, D. Ecological specialization and susceptibility to disturbance: conjectures and refutations. *The American naturalist* **159**, 606–623 (2002). |
| Darvill et al. † | Darvill, B., Knight, M. E. & Goulson, D. Use of genetic markers to quantify bumblebee foraging range and nest density. *Oikos* **107**, 471–478 (2004). |
| Quaranta et al. † | Quaranta, M. *et al.* Wild bees in agroecosystems and semi-natural landscapes. 1997–2000 collection period in Italy. *Bulletin of Insectology* **57**, 11–61 (2004). |
| Hanley (2005, unpublished data) † | Hanley, M.E. Unpublished data of bee diversity in UK croplands (2005) |
| Shuler et al. † | Shuler, R. E., Roulston, T. H. & Farris, G. E. Farming practices influence wild pollinator populations on squash and pumpkin. *Journal of Economic Entomology* **98**, 790–795 (2005). |
| Blanche et al. † | Blanche, K. R., Ludwig, J. A. & Cunningham, S. A. Proximity to rainforest enhances pollination and fruit set in orchards. *Journal of Applied Ecology* **43**, 1182–1187 (2006). |
| Diekötter et al. † | Diekötter, T., Walther-Hellwig, K., Conradi, M., Suter, M.& Frankl, R. Effects of landscape elements on the distribution of the rare bumblebee species Bombus muscorum in an agricultural landscape. *Biodiversity and Conservation* **15**, 57–68 (2006). |
| Marshall et al. † | Marshall, E. J. P., West, T. M. & Kleijn, D. Impacts of an agri-environment field margin prescription on the flora and fauna of arable farmland in different landscapes. *Agriculture, Ecosystems & Environment* **113**, 36–44 (2006). |
| McFrederick & LeBuhn †‡ | McFrederick, Q. S. & LeBuhn, G. Are urban parks refuges for bumble bees Bombus spp. (Hymenoptera: Apidae)?*Biological Conservation* **129**, 372–382 (2006). |
| Hatfield & Lebuhn † | Hatfield, R. & Lebuhn, G. Patch and landscape factors shape community assemblage of bumble bees, Bombusspp. (Hymenoptera: Apidae), in montane meadows. *Biological Conservation* **139**, 150–158 (2007). |
| Herrmann et al. †‡ | Herrmann, F., Westphal, C., Moritz, R. F. A. & Steffan-Dewenter, I. Genetic diversity and mass resources promote colony size and forager densities of a social bee (Bombus pascuorum) in agricultural landscapes. *Molecular Ecology* **16**, 1167–1178 (2007). |
| Meyer et al. † | Meyer, B., Gaebele, V. & Steffan-Dewenter, I. D. Patch size and landscape effects on pollinators and seed set of the Horseshoe Vetch, Hippocrepis comosa, in an agricultural landscape of Central Europe. *Entomologia Generalis* **30**, 173–185 (2007). |
| Öckinger & Smith | Öckinger, E. & Smith, H. G. Semi-natural grasslands as population sources for pollinating insects in agricultural landscapes. *Journal of Applied Ecology* **44**, 50–59 (2007). |
| Billeter et al. †, Diekötter et al. † and Le Féon et al. † | Billeter, R. *et al.* Indicators for biodiversity in agricultural landscapes: a pan-European study. *Journal of Applied Ecology* **45**, 141–150 (2008).  Diekötter, T., Billeter, R. & Crist, T. O. Effects of landscape connectivity on the spatial distribution of insect diversity in agricultural mosaic landscapes. *Basic and Applied Ecology* **9**, 298–307 (2008).  Le Féon, V. *et al.* Intensification of agriculture, landscape composition and wild bee communities: A large scale study in four European countries. *Agriculture, Ecosystems & Environment* **137**, 143–150 (2010). |
| Franzén & Nilsson † | Franzén, M. & Nilsson, S. G. How can we preserve and restore species richness of pollinating insects on agricultural land? *Ecography* **31**, 698–708 (2008). |
| Goulson et al. † | Goulson, D., Lye, G. C. & Darvill, B. Diet breadth, coexistence and rarity in bumblebees. *Biodiversity and Conservation* **17**, 3269–3288 (2008). |
| Kohler et al. † | Kohler, F., Verhulst, J., Van Klink, R. & Kleijn, D. At what spatial scale do high-quality habitats enhance the diversity of forbs and pollinators in intensively farmed landscapes? *Journal of Applied Ecology* **45**, 753–762 (2008). |
| Kwaiser & Hendrix | Kwaiser, K. S. & Hendrix, S. D. Diversity and abundance of bees (Hymenoptera: Apiformes) in native and ruderal grasslands of agriculturally dominated landscapes. *Agriculture, Ecosystems & Environment* **124**, 200–204 (2008). |
| Julier & Roulston † | Julier, H. E. & Roulston, T. H. Wild bee abundance and pollination service in cultivated pumpkins: Farm management, nesting behavior and landscape effects. *Journal of Economic Entomology* **102**, 563–573 (2009). |
| Knight et al. †‡ | Knight, M. E. *et al.* An interspecific comparison of foraging range and nest density of four bumblebee (Bombus) species. *Molecular Ecology* **14**, 1811–1820 (2005). |
| Vergara & Badano † | Vergara, C. H. & Badano, E. I. Pollinator diversity increases fruit production in Mexican coffee plantations: The importance of rustic management systems. *Agriculture, Ecosystems & Environment* **129**, 117–123 (2009). |
| Albrecht et al. | Albrecht, M. *et al.* Effects of ecological compensation meadows on arthropod diversity in adjacent intensively managed grassland. *Biological Conservation* **143**, 642–649 (2010). |
| Davis et al. †‡ | Davis, E. S., Murray, T. E., Fitzpatrick, Ú., Brown, M. J. F. & Paxton, R. J. Landscape effects on extremely fragmented populations of a rare solitary bee, *Colletes floralis. Molecular Ecology* **19**, 4922–4935 (2010). |
| Goulson et al. † | Goulson, D. *et al.* Effects of land use at a landscape scale on bumblebee nest density and survival. *Journal of Applied Ecology* **47**, 1207–1215 (2010). |
| Malone et al. †‡ | Malone, L. *et al.* Observations on bee species visiting white clover in New Zealand pastures. *Journal of Apicultural Research* **49**, 284–286 (2010). |
| Quintero et al. † | Quintero, C., Morales, C. L. & Aizen, M. A. Effects of anthropogenic habitat disturbance on local pollinator diversity and species turnover across a precipitation gradient. *Biodiversity and Conservation* **19**, 257–274 (2010). |
| Redpath et al. † | Redpath, N., Osgathorpe, L. M., Park, K. & Goulson, D. Crofting and bumblebee conservation: The impact of land management practices on bumblebee populations in northwest Scotland. *Biological Conservation* **143**, 492–500 (2010). |
| Bates et al. † | Bates, A. J. *et al.* Changing bee and hoverfly pollinator assemblages along an urban-rural gradient. *PLoS ONE* **6**, e23459 (2011). |
| Hanley et al. † | Hanley, M. E. *et al.* Increased bumblebee abundance along the margins of a mass flowering crop: evidence for pollinator spill-over. *Oikos* **120**, 1618–1624 (2011). |
| Blake et al. † | Blake, R. J., Westbury, D. B., Woodcock, B. A., Sutton, P. & Potts, S. G. Enhancing habitat to help the plight of the bumblebee. *Pest Management Science* **67**, 377–379 (2011). |
| Connop et al. †‡ | Connop, S., Hill, T., Steer, J. & Shaw, P. Microsatellite analysis reveals the spatial dynamics of Bombus humilisand Bombus sylvarum. *Insect Conservation and Diversity***4**, 212–221 (2011). |
| Hanley (unpublished data, 2011) † | Hanley, M.E. Unpublished data of bee diversity in UK croplands and urban habitats (2011). |
| Holzschuh et al. | Holzschuh, A., Dormann, C. F., Tscharntke, T. & Steffan-Dewenter, I. Expansion of mass-flowering crops leads to transient pollinator dilution and reduced wild plant pollination. *Proceedings of the Royal Society B: Biological Sciences* **278**, 3444–3451 (2011). |
| Power & Stout† | Power, E. F. & Stout, J. C. Organic dairy farming: impacts on insect-flower interaction networks and pollination. *Journal of Applied Ecology* **48**, 561–569 (2011). |
| Richards et al. † | Richards, M. *et al.* Bee diversity in naturalizing patches of Carolinian grasslands in southern Ontario, Canada. *The Canadian Entomologist* **143**, 279–299 (2011). |
| Samnegård et al. † | Samnegård, U., Persson, A. S. & Smith, H. G. Gardens benefit bees and enhance pollination in intensively managed farmland. *Biological Conservation* **144**, 2602–2606 (2011). |
| Schüepp et al. † | Schüepp, C., Herrmann, J. D., Herzog, F. & Schmidt-Entling, M. H. Differential effects of habitat isolation and landscape composition on wasps, bees, and their enemies. *Oecologia* **165**, 713–721 (2011). |
| Tonietto et al. † | Tonietto, R., Fant, J., Ascher, J., Ellis, K. & Larkin, D. A comparison of bee communities of Chicago green roofs, parks and prairies. *Landscape and Urban Planning* **103**, 102–108 (2011). |
| Weiner et al. | Weiner, C. N., Werner, M., Linsenmair, K. E. & Blüthgen, N. Land use intensity in grasslands: Changes in biodiversity, species composition and specialisation in flower visitor networks. *Basic and Applied Ecology* **12**, 292–299 (2011). |
| Fierro et al.[95](https://www.nature.com/articles/srep31153#ref95) †‡ | Fierro, M., Cruz-López, L., Sánchez, D., Villanueva-Gutiérrez, R. & Vandame, R. Effect of biotic factors on the spatial distribution of stingless bees (Hymenoptera: Apidae, Meliponini) in fragmented Neotropical habitats. *Neotropical Entomology* **41**, 95–104 (2012). |
| Lentini et al. † | Lentini, P. E., Martin, T. G., Gibbons, P., Fischer, J. & Cunningham, S. A. Supporting wild pollinators in a temperate agricultural landscape: Maintaining mosaics of natural features and production. *Biological Conservation***149**, 84–92 (2012). |
| Mudri-Stojnic et al. †‡ | Mudri-Stojnic, S., Andric, A., Józan, Z. & Vujic, A.Pollinator diversity (Hymenoptera and Diptera) in semi-natural habitats in Serbia during summer. *Archives of Biological Sciences* **64**, 777–786 (2012). |
| Osgathorpe et al.[133](https://www.nature.com/articles/srep31153#ref133) † | Osgathorpe, L. M., Park, K. & Goulson, D. The use of off-farm habitats by foraging bumblebees in agricultural landscapes: implications for conservation management. *Apidologie* **43**, 113–127 (2012). |
| Verboven et al.[97](https://www.nature.com/articles/srep31153#ref97) † | Verboven, H. A. F., Brys, R. & Hermy, M. Sex in the city: Reproductive success of Digitalis purpurea in a gradient from urban to rural sites. *Landscape and Urban Planning***106**, 158–164 (2012). |
| Yoon et al. | Yoon, H. J., Lee, K. Y., Kim, M. A. & Park, I. G. Local distribution and floral preferences of founder bumblebee queens in Korea. *Journal of Apiculture* **27**, 169–178 (2012). |
| Cunningham et al. † | Cunningham, S. A., Schellhorn, N. A., Marcora, A. & Batley, M. Movement and phenology of bees in a subtropical Australian agricultural landscape. *Austral Ecology* **38**, 456–464 (2013). |
| Grass et al. †‡ | Grass, I., Berens, D. G., Peter, F. & Farwig, N. Additive effects of exotic plant abundance and land-use intensity on plant-pollinator interactions. *Oecologia* **173**, 913–923 (2013). |
| Litchwark (unpublished thesis, 2013)† | Litchwark, S. A. Honeybee declines in a changing landscape: interactive effects of honeybee declines and land-use intensification on pollinator communities. *MSc Thesis*, University of Canterbury. |
| Rader et al. † | Rader, R., Bartomeus, I., Tylianakis, J. M. & Laliberté, E. The winners and losers of land use intensification: pollinator community disassembly is non-random and alters functional diversity. *Diversity and Distributions* **20**, 908–917 (2014). |
| Tylianakis et al. † | Tylianakis, J. M., Klein, A.-M. & Tscharntke, T. Spatiotemporal variation in the diversity of Hymenoptera across a tropical habitat gradient. *Ecology* **86**, 3296–3302 (2005). |
| Barlow et al. † | Barlow, J. *et al.* Quantifying the biodiversity value of tropical primary, secondary, and plantation forests. *Proceedings of the National Academy of Sciences* **104**, 18555–18560 (2007). |
| Meyer et al. † | Meyer, B., Jauker, F. & Steffan-Dewenter, I. Contrasting resource-dependent responses of hoverfly richness and density to landscape structure. *Basic and Applied Ecology***10**, 178–186 (2009).  Jauker, B., Krauss, J., Jauker, F. & Steffan-Dewenter, I. Linking life history traits to pollinator loss in fragmented calcareous grasslands. *Landscape Ecology* **28**, 107–120 (2013). |
| Schüepp et al. † | Schüepp, C., Rittiner, S. & Entling, M. H. High bee and wasp diversity in a heterogeneous tropical farming system compared to protected forest. *PLoS ONE* **7**, e52109 (2012). |
| Parra-H & Nates-Parra † | Parra-H, A. & Nates-Parra, G. Variation of the orchid bees community (Hymenoptera: Apidae) in three altered habitats of the Colombian “llano” piedmont. *Revista de biologia tropical* **55**, 931–941 (2007). |
| Smith-Pardo & Gonzalez † | Smith-Pardo, A. & Gonzalez, V. H. Diversidad de abejas (Hymenoptera: Apoidea) en estados sucesionales del bosque humedo tropical TT - Bee diversity (Hymenoptera: Apoidea) in a tropical rainforest succession. *Acta Biológica Colombiana* **12**, 43–55 (2007). |

**Appendix S2: Building complete phylogenetic trees**

The pastis package automatically defines constraints for the placement of missing species based on their taxonomic affinities, with congeners constrained to be sisters in a monophyletic clade, unless the phylogeny provides evidence against genus monophyly. Constraints can also be defined manually. These constraints are then used as an input to estimate phylogenetic trees using a Yule process in MrBayes (Ronquist et al. 2012). The resulting posterior distribution of trees represents a range of possible topologies, with all species included in the trees.

We used this procedure on twenty-eight bee clades with missing species (these were usually subfamilies or tribes which had greater than 85\% bootstrap support; See table S2.1 for details); this ensures that the backbone topology of the original tree is not altered, but placement of species within the clade will vary. For each clade, a sister taxon was constrained to be the outgroup. Phylogenetic trees were estimated using MrBayes version 3.2 (Ronquist et al. 2012) via CIPRES (Miller et al. 2010) for at least 100,000,000 generations and four runs, with samples taken every 10,000 generations. Tracer version 1.6 (Rambaut et al. 2013) was used to track effective sample sizes to assess convergence of parameter estimates. The standard deviations of split frequencies of the four independent runs were also assessed (Ronquist et al. 2012).

From each of the converged runs, we randomly (without replacement) sub-sampled from the post-burn-in (10% burn-in) posterior distribution of each bee clade to produce 1000 within-clade trees. A large clade of apid bees did not reach parameter convergence in any of the four independent runs, but measures of phylogenetic signal within the clade were not significantly different between runs (analysis of variance, F_3,998_ = 0.378, n.s.), so a random sample (without replacement) of 1,000 trees from all runs was taken. The original tree was rate smoothed using PATHd8 (Britton et al. 2007), with the root age constrained to one; incomplete clades were then pruned. To graft the completed within-clade trees back onto this pruned, rate-smoothed tree, the clade was first scaled to have a depth of 1. Following Jetz et al. (2012) we then rescaled the clade depth by dividing the depth of the ingroup node and the depth of the split between the clade and its outgroup, multiplied by the depth of the node linking the clade and its outgroup. The outgroup was dropped before the clade was grafted onto the backbone tree.

*Topological constraints*

For species without congeners in the phylogenetic tree, we used higher-level taxonomic constraints for species placement; we only used such constraints where nodes had greater than 95\% bootstrap support. Caenaugochlora, Chlerogella, Pereirapis, Pseudaugochlora, Chalepogenus and Agapostemonoides were restricted to their respective tribes, where monophyly was strongly supported (Hedtke et al. 2013). Note that Agapostemonoides was constrained within the tribe Caenohalictini, which is sometimes considered only a subtribe within the Halictini tribe (Danforth et al. 2008). Pachyprosopis (Euryglossinae: Colletidae) was constrained to be sister to the tribes Euryglossinae, Scrapterinae, and Xeromelissinae, but species were not permitted to enter the clades formed by the Xeromelissinae or Hylaeinae (Almeida and Danforth 2009, Hedtke et al. 2013). The genus Ceylalictus was constrained to be placed within its subfamily, Nomioidinae. Where synonyms were identified by a taxonomic expert (MK), these were merged.

Where the published phylogeny had dubious species placements (i.e., the placement of Ceratina japonica and Anthophora pillipes outside of their otherwise monophyletic groups and placed with fairly distantly related species) that were noted as such by the authors of the tree (Hedtke et al. 2013), these were considered missing species and their placement was constrained automatically using the pastis package.

Table S2.1: Details of within-clade trees run using MrBayes to include missing species (using the Birth-Death polytomy resolver). The species tree is the original phylogenetic tree produced by Hedtke et al. (2013) and used throughout the paper. The genus tree from Hedtke et al (2013) was also used to explore phylogenetic relationships among genera.

| **Family** | **Clade** | **Bootstrap support (Genus tree)** | **Bootstrap support (Species tree)** | **Node in species tree** | **Number of generations (millions)** |
| --- | --- | --- | --- | --- | --- |
| Apidae | Meloponini, Bombini, Apini | 74 | 91 | 2300 | 400 |
| Apidae | Euglossini | 100 | 100 | 2296 | 100 |
| Apidae | Centridini (Centris) | 100 | 100 | 2287 | 100 |
| Apidae | Centridini (Epicharis) | 100 | 100 | 2284 | 100 |
| Apidae | Allodapini and Ceratinini | 100 |  | 2211 | 100 |
| Apidae | Manuelini |  |  | manuelini | 100 |
| Apidae | Xylocopini |  | 85 | 2236 | 200 |
| Apidae | Epeolini, Brachynomadini, ammobatoidini, Biastini, Townsendiellini, Neolarrini, Nomadini, Hexepeoplini, Ammobatini, Melectini, Caenoprospidini, Ericrocidini, Rhathymini, Esepeolini, Osirini (Epeoloides), Protepeolini, Osirini (Osiris), Tetrapedini (Coelioxoides), Osirini (Parepeolus) | 99 | 100 | 2088 | 400 |
| Apidae | Eucerini, Ancylini, Emphorini, Tapinotaspidini, Exomalopsini, Emphorini | 100 | 100 | 2154 | 400 |
| Apidae | Anthophorini | 100 | 100 | 2081 | 100 |
| Andrenidae | Andreninae | 100 | 100 | 1457 | 400 |
| Andrenidae | Panurginae | 99 | 99 | 1465 | 100 |
| Halictidae | Halictini | 97 |  | 1774 | 400 |
| Halictidae | Sphecodini | 100 |  | 1762 | 100 |
| Halictidae | Caenohalictini | 99 |  | 1749 | 100 |
| Halictidae | Augochlorini | 100 |  | 1722 | 200 |
| Halictidae | Nomiinae | 100 |  | 1707 | 100 |
| Halictidae | Rophitinae | 100 |  | 1687 | 100 |
| Halictidae | Nomiodinae | 100 |  | 1719 | 100 |
| Melittidae | Melittidae | 98 |  | 1412 | 100 |
| Colletidae | Hylaeinae, Scrapterinae, Euryglossinae, Xeromelissinae | 97 |  | 1605 | 400 |
| Colletidae | Colletini | 100 |  | 1590 | 100 |
| Colletidae | Neophashaeinae | 100 | 90 | 1535 | 200 |
| Megachilidae | Megachilini | 100 | 100 | 1933 | 200 |
| Megachilidae | Lithurgini | 100 | 100 | 1880 | 100 |
| Megachilidae | Heriades | 100 | 100 | 2000 | 100 |
| Megachilidae | Anthidini (anthidiellum and anthidini) | 100 | 100 | 1900 | 200 |
| Megachilidae | Osmini | 100 | 100 | 2008 | 100 |

**Appendix 3: Functional Traits**

Table 3.1: Original and coarsened factor levels of species traits

| Trait | Coarsened factor levels | Original factor levels | Rationale |
| --- | --- | --- | --- |
| Nesting trait | Excavators | Excavators in the soil or vegetation | This trait was coarsened to represent two distinct nesting strategies: those that build their own holes versus those that don't. Excavators are particular about nesting sites, often requiring hard, bare ground or pithy stems, whilst those that don't excavate use existing cavities or old nesting sites, regardless of nest location. |
|  | Non-excavators | Carder bees, renters, masons, cleptoparasites and social parasites |  |
| Sociality | Obligately solitary | Solitary, solitary or communal, communal, cleptoparasitic | The sociality of the species was defined according to how their offspring are raised, because this relates to reproductive capacity. Social species, or those that raise their young in social nests (such as social parasites), are able to produce greater numbers of offspring because there are more workers to provision those offspring. Primitively eusocial species are able to adjust their reproductive capacity, often according to resource requirements: for example, *Halictus rubicundus* is social in warmer, more resource rich areas but solitary in other areas. |
|  | Not obligately solitary | Highly eusocial, primitively eusocial, solitary/primitively eusocial, polymorphic, social parasites |  |
| Lecty | No Lecty status | No Lecty status | Species with no lecty status are those which do not collect their own pollen, for example cleptoparasites. Phenotypic flexibility can be considered as a form of generalism so species that can be either oligolectic or polylectic are considered in the same category as the pollen generalists. |
|  | Obligately oligolectic | Oligolectic |  |
|  | Polylectic/Flexible | Polylectic, oligolectic or polylactic |  |
| Voltinism | Univoltine | Univoltine | Species were split into two categories: those with only one generation per year, and those that do have or can have more than one generation per year, as the latter are predicted to be less impacted by local threats. |
|  | Multivoltine/Flexible | Bivoltine, multivoltine, univoltine or bivoltine, univoltine or multivoltine |  |

Table 3.2: Phylogenetic signal of bee species traits using both the incomplete phylogeny (Hedtke et al. 2013) and the gap-filled phylogeny.

| **Trait** | **Signal in the rate-smoothed tree** | **Signal in the completed trees (mean ± standard deviation)** |
| --- | --- | --- |
| Flight Season duration | λ = 0, n.s. | λ = 0.199 (± 0.29), p < 0.05 in 1000 trees |
| ITD | λ = 0.94, p < 0.001 | λ = 0.934 (± 0.01), p < 0.05 in 1000 trees |
| Nesting strategy | D = -0.39, p < 0.001 | D = -0.488 (± 0.04), p < 0.05 in 1000 trees |
| Sociality | D = -0.32, p < 0.001 | D = -0.321 (± 0.06), p < 0.05 in 1000 trees |
| Tongue length | D = -0.71, p < 0.001 | D = -0.687 (± 0.04), p < 0.05 in 1000 trees |
| Voltinism | D = 0.27, p < 0.05 | D = 0.618(± 0.09), p < 0.05 in 994 trees |
| Lecty status | λ = 0, n.s. | λ = 0.44 (± 0.5), p < 0.05 in 0 trees |

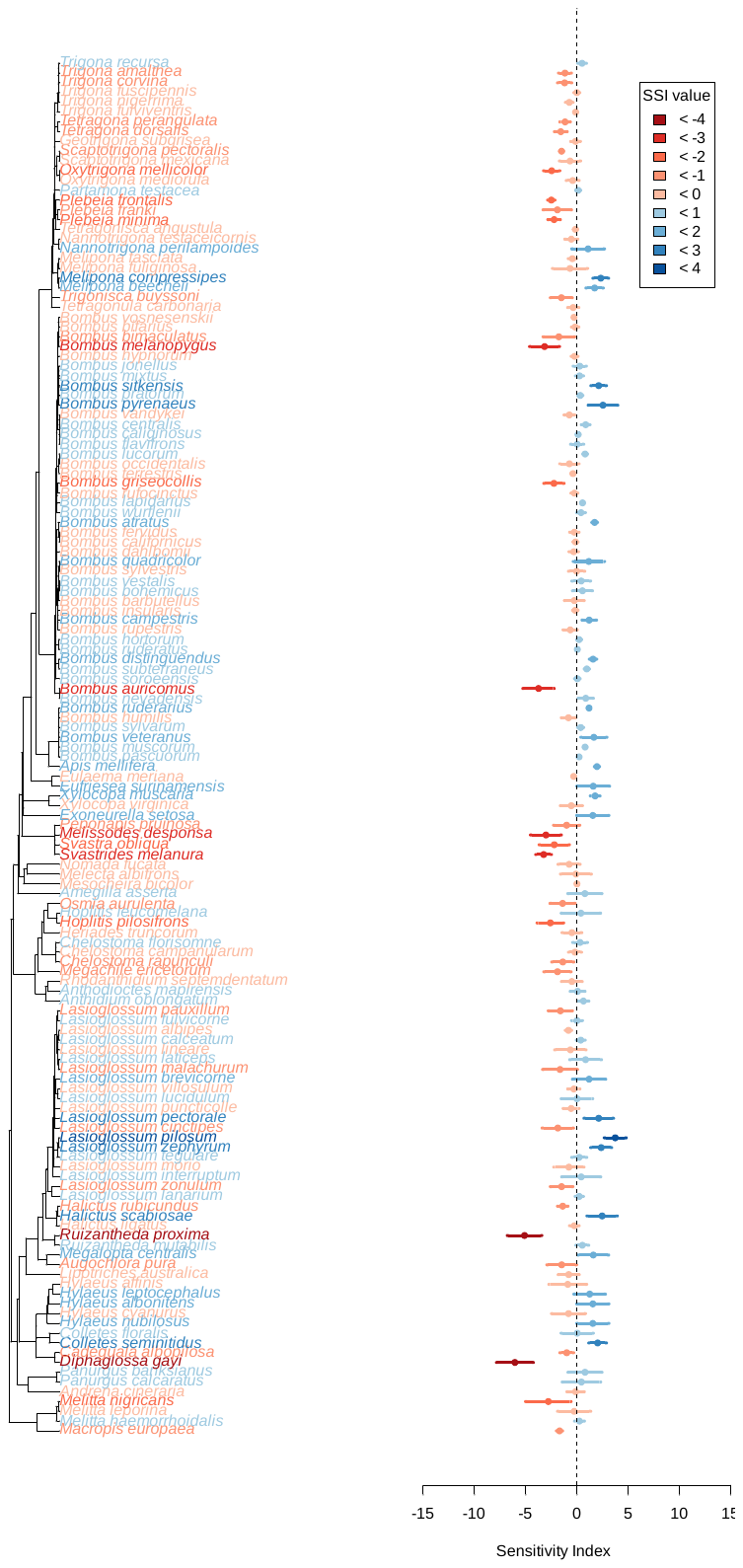
**Appendix 4: Distribution of species sensitivities across the rate-smoothed (incomplete) phylogeny**

Figure S4.1: Sensitivity of species to human-dominated land across the phylogeny. Tips of the phylogeny are coloured according to the species’ sensitivity: from blue to red indicated less to more sensitive. The right panel shows species sensitivity ± the standard error.

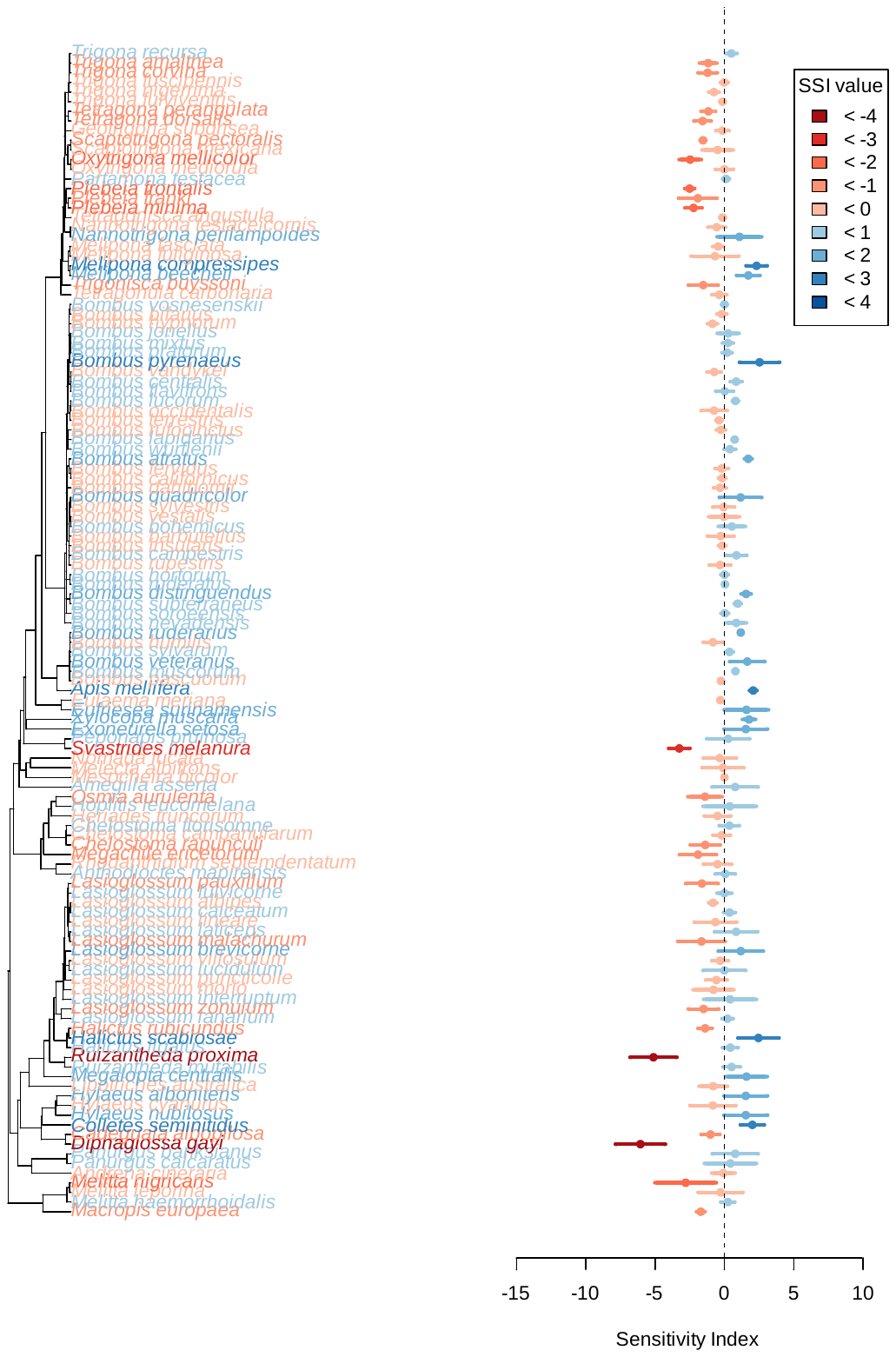
Figure S4.2: Sensitivity of species to agricultural land across the phylogeny. Tips of the phylogeny are coloured according to the species' sensitivity: from blue to red indicate less to more sensitive. The right panel shows species sensitivity ± the standard error.

Figure S4.3: Sensitivity of species to urban land across the phylogeny. Tips of the phylogeny are coloured according to the species' sensitivity: from blue to red indicate less to more sensitive. The right panel shows species sensitivity ± the standard error.

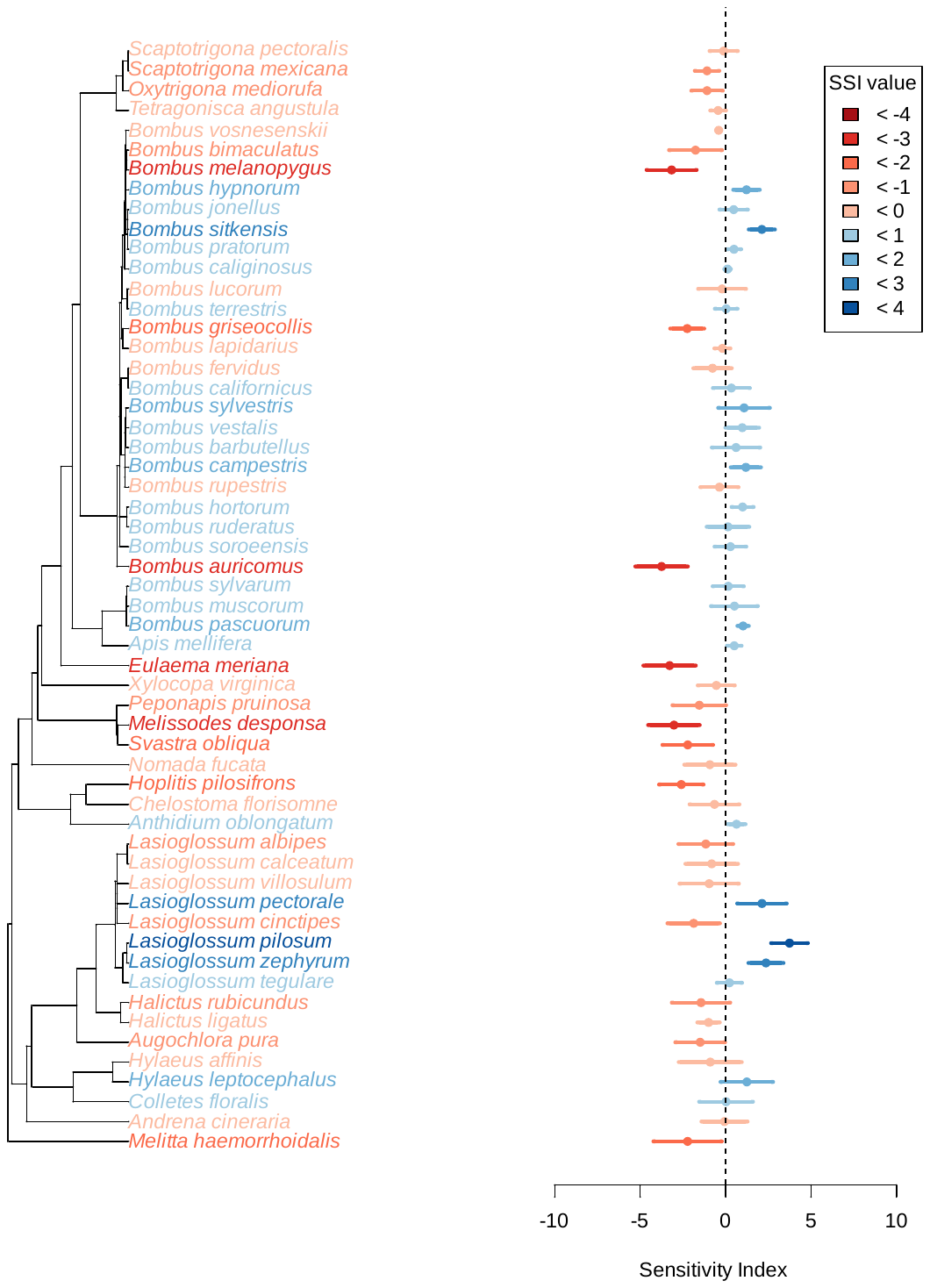

Figure S4.4: Sensitivity of species to agricultural intensification across the phylogeny. Tips of the phylogeny are coloured according to the species' sensitivity: from blue to red indicate less to more sensitive. The right panel shows species sensitivity ± the standard error.
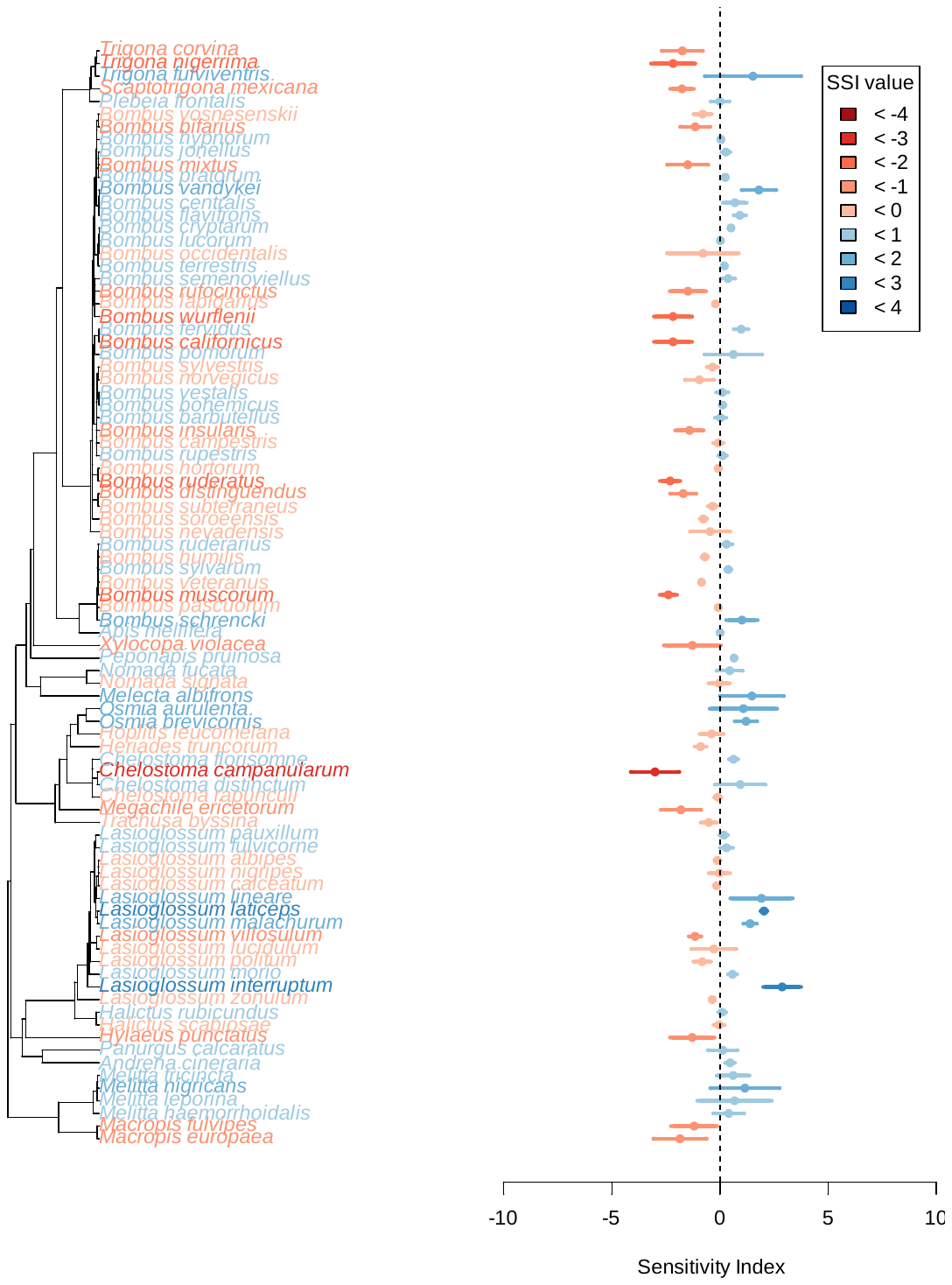

**Appendix 5: Relative Importance of Traits and Phylogeny**

Figure S5.1: Explanatory power R^2^ of models assessing species sensitivities as a function of alpha value, where a = 0 uses only functional traits to calculate the distance, a = 1 uses only phylogeny to calculate the distance, and a = 0.5 considers both traits and phylogeny equally. Model performance was also calculated where distances were randomly computed. Red triangles indicate the values for the rate-smoothed tree, where some species are missing from the tree. Numbers in red show the number of species included in these tests. Boxplots show the distribution of values across the 1,000 completed trees, with numbers in black showing the number of species included in these tests.

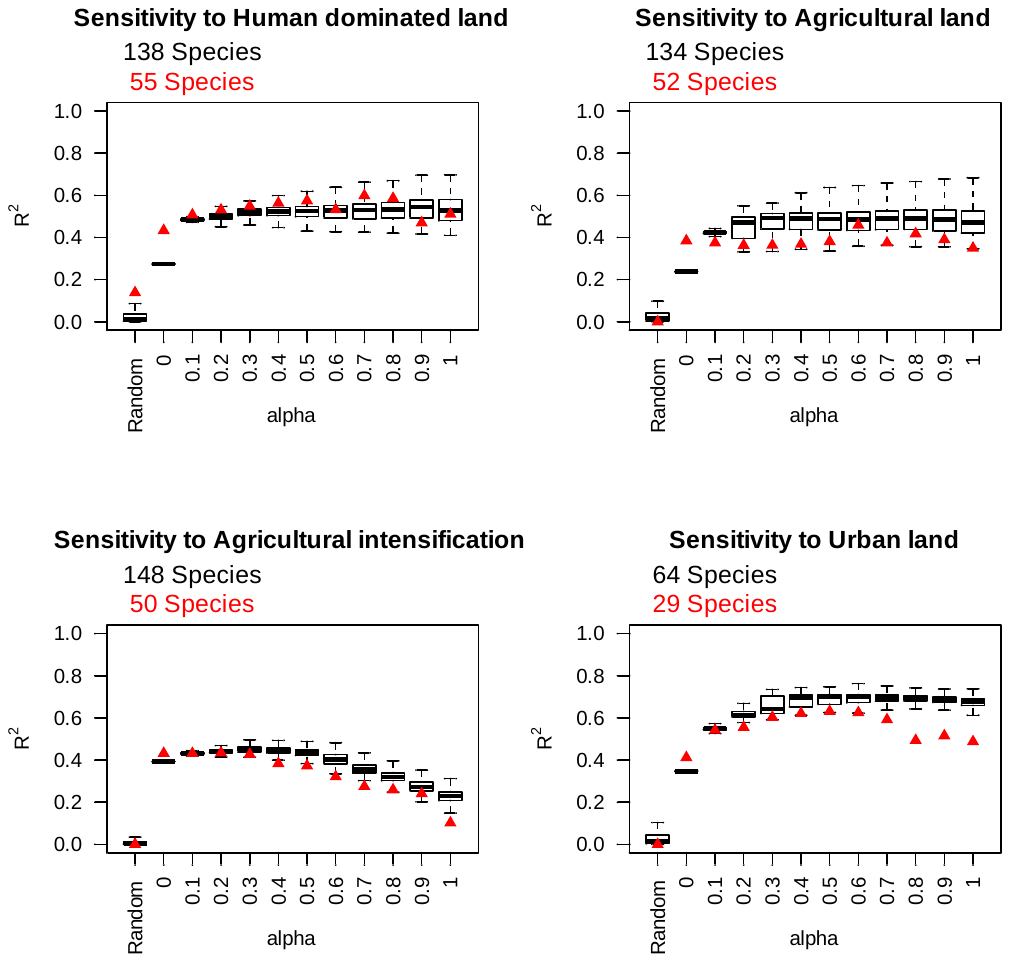

**Appendix 6: Representativeness of species included in the analysis**

The species present in local communities are likely to be non-random subsets of the phylogeny (Vamosi et al. 2009, Cavender-Bares et al. 2009). Using the incomplete phylogenetic tree (Hedtke et al. 2013), we assessed the phylogenetic diversity of species present in our dataset against a random subset of species from the phylogeny (999 times, using an independent swap algorithm to sample species, maintaining species occurrence frequency and species richness, ses.pd function, picante package: Kembel et al. 2010). The phylogenetic diversity of our sample was 16.03, compared to a mean phylogenetic diversity of null communities of 20.39 (standardised effect size of PD vs null communities = -4.13, *p* < 0.01).
